# Neuronal C-Reactive Protein Mediates Neuropathic Pain by Activating Nociceptive FcγRI-Coupled Signaling

**DOI:** 10.1101/2022.08.30.505953

**Authors:** Fan Liu, Si Su, Li Zhang, Yehong Fang, Huan Cui, Jianru Sun, Yikuan Xie, Chao Ma

**Affiliations:** National Human Brain Bank for Development and Function, Institute of Basic Medical Sciences Chinese Academy of Medical Sciences, School of Basic Medicine Peking Union Medical College, Beijing 100005, China; Department of Human Anatomy, Histology and Embryology, Neuroscience Center, Institute of Basic Medical Sciences Chinese Academy of Medical Sciences, School of Basic Medicine Peking Union Medical College, Beijing 100005, China; Department of Anesthesiology, Beijing Friendship Hospital, Capital Medical University, Beijing 100050, China; Chinese Institute for Brain Research, Beijing 102206, China

**Keywords:** Nociceptive neuron, IgG, Spleen tyrosine kinase, CD64, Chronic pain

## Abstract

**Background:** Neuropathic pain is difficult to treat in clinical practice, and the underlying mechanisms are insufficiently elucidated. Previous studies have demonstrated that Fcγ receptor I (FcγRI) is expressed in the neurons of the dorsal root ganglion (DRG) and may be involved in chronic pain.

**Methods:** Chronic constriction injury (CCI) was used to induce neuropathic pain in rats. Primary neuron-specific *Fcgr1* conditional knockout (CKO) rats were established by crossing rats carrying a *Fcgr1*^*loxP+/+*^ with the *Pirt*^*CRE*+^ line. Behavioral and molecular studies were conducted to evaluate the differences between wild-type and CKO rats after CCI.

**Results:** We first revealed that CCI activated neuronal FcγRI-related signaling in the DRG. CCI-induced neuropathic pain was alleviated in CKO rats. C-reactive protein (CRP) was increased in the DRG after nerve injury. Intraganglionic injection or overexpression of the recombinant CRP protein in the DRG evoked pain accompanied and activated neuronal FcγRI. CRP-evoked pain was significantly reduced in CKO rats. Furthermore, microinjection of native IgG into the DRG alleviated neuropathic pain and the activation of neuronal FcγRI-related signaling.

**Conclusions:** Our results indicate that the activation of neuronal CRP/FcγRI-related signaling plays an important role in the development of pain in CCI. Our findings may provide novel insights into the neuroimmune responses after peripheral nerve injury and might suggest potential therapeutic targets for neuropathic pain.

## 1. Introduction

Neuropathic pain is the most common form of chronic pain resulting from damage or disease of the somatosensory system, symptoms of which include spontaneous pain, hyperalgesia, and allodynia ^1^. Neuropathic pain, one of the main problems troubling human health, not only seriously affects patients’ quality of life but also places a huge financial burden on society. A systematic review of epidemiological studies reported that the prevalence of neuropathic pain has been estimated at 6.9-10.0% within the general population, and the number of neuropathic pain patients is increasing year by year ^2^. Because the pathogenesis of neuropathic pain is still unclear, the analgesic effects of existing treatments and drugs are not satisfactory and are often coupled with intolerable side effects. Therefore, it is urgent to study and elucidate the occurrence mechanism and pathogenesis of neuropathic pain and to find new therapeutic strategies and drug targets on this basis.

In previous research reports, the immune system and peripheral nervous system play an important role in neuropathic pain. Research has found that neuropathic pain is characterized by immune diseases ^3-5^. Peripheral nerve injury often results in the disintegration of nerve fibers, induces inflammation, and may release autoantigens ^6, 7^. In recent years, it has been reported that peripheral nociceptive neurons have immune sensing functions and immune recognition through the expression of immune-related receptors such as Fc-gamma receptors, Fc-epsilon receptors and Toll-like receptors ^8-13^. Our previous research revealed that partial neurons of the dorsal root ganglion (DRG) and trigeminal ganglion (TG) in rodents express FcγRI (a type of receptor with high affinity activation for IgG) and can be activated by IgG-IC to lead to pain and hyperalgesia ^12-14^. DRG neurons not only expressed FcγRI (coexpressed with markers of nociceptors such as TRPV1, IB4, CGRP and SP) but could also be activated by IgG-IC in vitro or in vivo ^12-14^. However, whether neuronal FcγRI in the DRG, which is activated by ligands, contributes to neuropathic pain has not been fully elucidated.

In the present study, we explored the potential roles of neuronal FcγRI in the DRG after nerve injury. We hypothesize that FcγRI-related signals in DRG neurons play a role in the development of neuropathic pain after nerve injury. In this study, we tested this hypothesis using an *Fcgr1* conditional knockout rat model of chronic constriction injury (CCI) of the sciatic nerves.

## 2. Methods

### 2.1. Animals

Male Sprague–Dawley rats were purchased from the National Institutes for Food and Drug Control (China). *Pirt*^Cre/+^;*Fcgr1*^*loxP/loxP*^ (*Fcgr1* CKO, maintained on a Sprague–Dawley genetic background) rats were described previously ^10^. Littermate rats were generated by interbreeding heterozygotes on the Sprague–Dawley genetic background. Male rats weighing 150-180 g were used, and rats were raised in standard cages (five per cage) in a 12-hour light/dark cycle and climate-controlled room with a specific pathogen-free environment. Animals were randomly assigned to treatment or control groups. All animal procedures performed in this study were reviewed and approved by the Institutional Animal Care and Use Committee of the Institute of Basic Medical Sciences, Chinese Academy of Medical Sciences, Peking Union Medical College (Beijing, China) and were conducted in accordance with the guidelines of the International Association for the Study of Pain.

### 2.2. Drugs and drug administration

We purchased recombinant CRP protein (Cat: 80041-R08H) from Sino Biological Inc.; ovalbumin (OVA, Cat: A5503) from Sigma-Aldrich; rat anti-OVA IgG (Cat: ACMOV111R) from Agro-Bio; highly selective Syk inhibitor PRT062607 (P505-15) HCl (Cat: S8032) from Selleck; native rat gamma globulin (native IgG, Cat: 012-000-002, Jackson ImmunoResearch) obtained from healthy rats. AAV2/9-hSyn-CRP-ZsGreen and AAV2/9-hSyn-ZsGreen (control vector) were made by Hanbio (China). As previously described ^12, 14^, IgG-IC was prepared by using OVA as the antigen and rat anti-OVA IgG as the antibody. CRP protein was dissolved in 20 μl phosphate buffered saline (PBS). Rats were randomly divided into groups and injected intradermally into the plantar skin of the hindpaw with a 20 μl volume of CRP, IgG-IC or PBS (Vehicle).

### 2.3. Intraganglionic injection of rat DRG

As previously described ^15^, rats were anesthetized with pentobarbital sodium (40 mg/kg i.p.). Exposing the L_4_-L_5_ spinal nerve, a microinjection syringe was inserted into the L_4_-L_5_ spinal nerve under the epineurium until the tip reached the center of the DRG. Then, the drugs (5 μL of AAV vectors consisting of 10^12^ viral particles or native IgG) were delivered to the L_4_-L_5_ DRGs.

### 2.4. Rat model of neuropathic pain

We obtained CCI rats according to the Bennett and Xie model ^16^. We used sodium pentobarbital (40 mg/kg i.p.) to anesthetize rats and utilized 4-0 surgical catgut to loosely tie 4 ligatures around the sciatic nerve of the mid-thigh level on the right side with approximately 1 mm space between the knots. Rats in the sham group only received sciatic nerve exposure without ligation.

### 2.5. Behavioral tests of pain

As previously described ^17, 18^, mechanical and thermal hyperalgesia in rats was quantified. Experimental rats were randomly divided into groups, and researchers were blinded regarding the group allocation. Briefly, rats were placed in individual plastic cages (10 × 20 × 20 cm) on a mesh floor and allowed to acclimate for 30 min. Rats were acclimatized for three consecutive days before the behavioral test, and three measurements were averaged for each rat at 5-10 min intervals. The paw withdrawal threshold in response to mechanical stimuli was used to assess mechanical allodynia. The probe of an electric von Frey anesthesia meter (IITC Life Science) was applied perpendicularly to the hindpaw with no acceleration at a force. Acute withdrawal of the hindpaw was considered a positive response. Paw withdrawal latency in response to radiant heat was used to evaluate thermal hyperalgesia. The radiant heat source of a thermal stimulator (BME-410C Plantar Test Apparatus) was focused on the hindpaw plantar surface of rats, which were positioned on the floor of the cage. When the rat moved or licked the hindpaw, the thermal stimulus was terminated, and the time from initiation to termination was recorded. A maximum cut-off time of 20 s was applied to prevent tissue damage in the thermal stimulus tests.

### 2.6. Immunoprecipitation and immunoblotting

L_4_-L_5_ DRGs were harvested from rats and snap-frozen in liquid nitrogen. Total proteins were extracted with RIPA lysis buffer (Thermo Fisher Scientific) containing NP-40, protease inhibitors (CoWin Biosciences) and phosphatase inhibitors (CoWin Biosciences). According to the protocol of the Pierce Crosslink Magnetic IP/Co-IP Kit (Fisher Scientific, #88805), DRG tissue lysates were immunoprecipitated over 2 hours at 4°C with protein A/G coupled with antibodies (rabbit anti-FcγRI 1:1000, Cat: 80016-R015, Sino Biological Inc.; rabbit anti-Syk 1:1000, Cat: ab40781, Abcam or control negative antibody (Cat: 3900, Cell Signaling Technology)). Proteins were resolved by 10% SDS–PAGE, transferred onto PVDF membranes (Merck Millipore), and immunoblotted with primary antibodies.

As previously described ^17, 18^, all homogenized samples of lumbar DRGs were centrifuged, and supernatants were mixed with SDS–PAGE loading buffer for 5 minutes at 95 °C. Thirty micrograms of each protein sample was separated by 10% SDS–PAGE and then transferred to PVDF membranes. All membranes were blocked with 5% BSA in TBST for 1 hour at room temperature and subsequently incubated with rabbit anti-FcγRI (1:1000, Cat: 80016-R015, Sino Biological Inc.), rabbit anti-FcRγ (1:500, Cat: ab151986, Abcam), rabbit anti-Syk (1:1000, Cat: ab40781, Abcam), rabbit anti-pSyk (1:1000, Cat: PA5-36692, Thermo Fisher Scientific), rabbit anti-pSrc (1:1000, Cat: ab185617, Abcam), rabbit anti-Src (1:1000, Cat: ab109381, Abcam), rabbit anti-CRP (1:500, Cat: ab259862, Abcam), rabbit anti-Vav1 (1:500, Cat: D155205, Sangon Biotech), mouse anti-pNF-κB p65 (1:500, Cat: 13346, Cell Signaling Tech), rabbit anti-NLRP3 (1:1000, Cat: ab263899, Abcam), rabbit anti-IL-18 (1:1000, Cat: PAB16177, Abnova), rabbit anti-IL-1β (1:1000, Cat: ab254360, Abcam) or mouse anti-GAPDH (1:2000, Cat: ab8245, Abcam) primary antibodies. The corresponding secondary antibodies (HRP-conjugated goat anti-rabbit or goat anti-mouse, 1:5000, CoWin Biosciences) were probed for 1 h at room temperature. The results were detected with an enhanced chemiluminescence reagent eECL Kit (Cat: CW0049, CoWin Biosciences).

### 2.7. Immunofluorescence staining

Fresh 4% paraformaldehyde was perfused into rats deeply anesthetized with sodium pentobarbital (40 mg/kg) through the ascending aorta. The L_4_-L_5_ DRGs and spinal dorsal horn were harvested, postfixed in 4% paraformaldehyde for 4 h and then dehydrated in 30% sucrose overnight at 4°C. The frozen tissue was sectioned to a thickness of 12 μm in a cryostat. After permeabilization with 0.2% Triton X-100 in PBS for 15 min and incubation with blocking buffer (10% normal donkey serum) for 1 h, the tissue sections were incubated with primary antibodies, such as rabbit anti-FcγRI (1:500, Cat: 80016-R015, Sino Biological Inc.), goat anti-FcRγ (1:100, Cat: sc-33496, Santa Cruz Biotech), rabbit anti-pSyk (1:100, Cat: PA5-36692, Thermo Fisher Scientific), rabbit anti-Syk (1:100, Cat: ab40781, Abcam), rabbit anti-Src (1:100, Cat: ab109381, Abcam), rabbit anti-CRP (1:200, Cat: ab259862, Abcam), rabbit anti-Vav1 (1:100, Cat: D155205, Sangon Biotech), rabbit anti-Sos1 (1:200, Cat: ab140621, Abcam), mouse anti-pNF-κB p65 (1:100, Cat: 13346, Cell Signaling Tech), rabbit anti-NLRP3 (1:200, Cat: ab263899, Abcam), rabbit anti-IL-18 (1:100, Cat: PAB16177, Abnova), rabbit anti-IL-1β (1:100, Cat: ab254360, Abcam), guinea pig anti-TRPV1 (1:800, Cat: ab10295, Abcam) and guinea pig anti-PGP9.5 (1:200, Cat: ab10410, Abcam) in PBS with 10% normal donkey serum overnight at 4°C. Afterwards, the slides were incubated with the proper secondary antibodies (Alexa Fluor 594-conjugated donkey anti-rabbit, 1:600; Alexa Fluor 594-conjugated donkey anti-guinea pig, 1:600; Alexa Fluor 488-conjugated donkey anti-mouse, 1:600 and Alexa Fluor 488-conjugated donkey anti-guinea pig, 1:600, Jackson ImmunoResearch) or Alexa Fluor 488-conjugated IB4 (1:200, Cat: I21411, Thermo Fisher Scientific) for 1 h. Slides were then washed in PBS and cover-slipped with VECTASHIELD Mounting Medium with DAPI (Cat: H-1200, Vector lab). Images were captured by a microscopic imaging system (Olympus BX61 and FluoView software), and the percentages of positive neurons were calculated and statistically analysed.

### 2.8. RNA sequencing

As previously described ^10^, L_4_ to L_5_ lumbar DRGs were obtained from 4 rats in the sham group (15 days after operation), 4 rats in the CCI+Veh group (15 days after operation) and 4 rats in the CCI+IgG group (15 days after operation). Total RNA was extracted using TRIzol reagent (Thermo Fisher Scientific). RNA degradation and contamination were monitored on 1% agarose gels. RNA integrity was assessed using the RNA Nano 6000 Assay Kit of the Agilent Bioanalyzer 2100 system (Agilent Technologies, USA). Sequencing libraries were generated using the NEBNext UltraTM RNA Library Prep Kit for Illumina (NEB, USA) following the manufacturer’s recommendations. The clustering of the index-coded samples was performed on a cBot Cluster Generation System using TruSeq PE Cluster Kit v4-cBot-HS (Illumina) according to the manufacturer’s instructions. After cluster generation, the library preparations were sequenced on an Illumina HiSeq 4000 platform, and paired-end 150 bp reads were generated. Raw data (raw reads) in fastq format were first processed through in-house Perl scripts. In this step, clean data (clean reads) were obtained by removing reads containing adapters, reads containing poly-N sequences and low-quality reads from the raw data. At the same time, the Q20, Q30, GC content and sequence duplication level of the clean data were calculated. All downstream analyses were based on clean data with high quality. Gene function was annotated based on the following databases: Nt (NCBI nonredundant nucleotide sequences); KO (KEGG Orthology database); and GO (Gene Ontology). Quantification of gene expression levels was estimated by fragments per kilobase of transcript per million fragments mapped.

### 2.9. Statistical analysis

Data values are expressed as the group mean ± SEM. Statistical analyses were performed using SPSS software (version 17.0). Student’s t test was used to analyse the statistical significance of differences between 2 groups. One-way analysis of variance (ANOVA) followed by Scheffe’s post-hoc test was used to determine statistical comparisons of differences among 3 or more groups. Two-way ANOVA followed by the Bonferroni post hoc test was used to determine significant differences in pain behavior. *p* < 0.05 was considered statistically significant. Differential expression analysis of the 2 groups was performed using the DESeq R package. Genes with an adjusted *p* value of less than 0.05 according to the DESeq analysis were defined as differentially expressed.

## 3. Results

### 3.1. Neuronal FcγRI is upregulated in the DRG after peripheral nerve injury

Previous studies demonstrated that neuronal FcγRI in the DRG could be directed by IgG-IC and mediate joint pain in animal models of RA ^13, 14, 19^. Peripheral nerve injury-induced protein changes in DRGs play a critical role in neuropathic pain generation. Chronic constriction injury of the sciatic nerve is a neuropathic pain model ^16^. First, we examined the protein expression of FcγRI and FcRγ in the nerve-injured DRGs of CCI rats. Our western blot analysis showed that CCI induced a long-lasting increase in FcγRI and FcRγ protein levels in the nerve-injured DRGs of rats (Figure 1a). To determine the cellular distribution of FcγRI and FcRγ, we then performed double immunolabelling using PGP9.5 (neuronal marker) with FcγRI or FcRγ. Immunostaining showed that FcγRI and FcRγ were predominantly immunoreactive in DRG neurons (Figure 1b, c).

**Figure 1.**
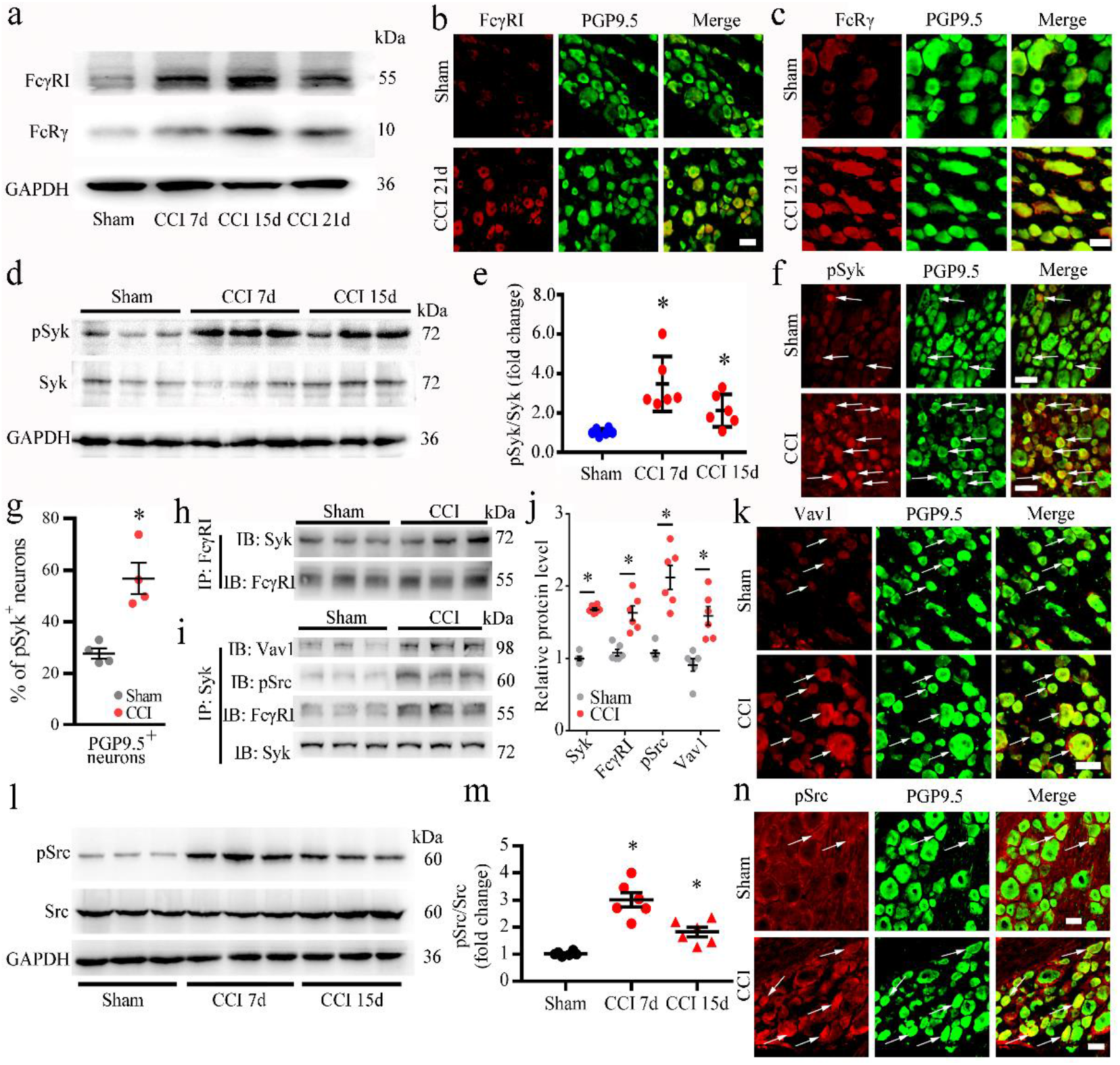
Expression, activation, and cellular distributions of FcγRI/Syk-related downstream signaling proteins in rat DRGs after CCI surgery. **(a)** Western blot analysis of the protein levels of FcγRI and FcRγ (b) in DRGs from sham, CCI 7 d, CCI 15 d and 21 d rats after surgery. **(b)** Double immunofluorescence showing the cellular distribution of FcγRI (red) and the neuronal marker PGP9.5 (green) in DRGs from sham and CCI 21 d rats after surgery. **(c)** Double immunofluorescence showing the cellular distribution of FcRγ (red) with PGP9.5 (green) in DRGs from sham and 21 d rats after surgery. **(d-e)** Western blot analysis and quantification of the protein levels of pSyk in DRGs from sham, CCI 7 d and 15 d rats after surgery. **(f)** Double immunofluorescence showing the cellular distribution of pSyk (red) with the neuronal marker PGP9.5 (green) in DRGs from sham and CCI rats 15 d after surgery. **(g)** Proportion of pSyk-positive neurons in the DRGs of sham and CCI rats 15 d after surgery. \**p* < 0.05 versus sham, t test, n = 4/group. **(h)** Co-IP showed the interaction between FcγRI and Syk in DRG tissue from sham and CCI rats 15 d after surgery. **(i)** Co-IP showed the interaction between Syk and Vav1, pSrc and FcγRI in DRG tissue from sham and CCI rats 15 d after surgery. **(j)** Quantification of Co-IP of Syk, FcγRI, pSrc, and Vav1 in DRGs from sham and CCI rats 15 d after surgery. n = 6 rats/group, \**p* < 0.05 versus the sham group, one-way ANOVA. **(k)** Double immunofluorescence showing the cellular distribution of Vav1 in DRGs from sham and CCI rats 15 d after surgery. **(l-m)** Western blot analysis (l) and quantification (m) of the protein levels of pSrc and Src in DRGs from sham, CCI 7 d and 15 d rats after surgery. **(n)** Double immunofluorescence showing the cellular distribution of pSrc (red) with the neuronal marker PGP9.5 (green) in DRGs from sham and CCI rats 15 d after surgery. Scale bars: 50 μm in figure b, c, k and n; 30 μm in figure d and f, n = 4 rats/group. In figure a, d, e, i and m, n = 6 rats/group, \**p* < 0.05, \*\**p* < 0.01 versus the sham group, one-way ANOVA.

### 3.2. Neuronal Syk signaling is activated in the DRG after peripheral nerve injury

To determine whether increased FcγRI and FcRγ expression levels are associated with increased related signaling activity, we measured the level of Syk, which is coupled with FcγRI ^20, 21^. Syk is a member of the family of nonreceptor-type Tyr protein kinases ^21^. It, which is widely expressed in immune cells, is involved in coupling activated immunoreceptors such as FcγRs and FcεRI to downstream signaling events that mediate receptor activation ^21^. Syk is required for FcγRI-induced excitation of sensory neurons in the DRG ^22^. Therefore, we examined the expression, activation, and cellular distribution of Syk in the DRG after CCI surgery. The experimental results indicated that phosphorylated Syk (pSyk) was expressed at higher levels than sham surgery in the DRGs of CCI rats (Figure 1d, e). The total Syk protein level of the rat DRGs was not changed after CCI surgery (Figure 1d). Double immunofluorescence images showed that pSyk was expressed in the small neurons of the DRG and was significantly higher in DRG neurons after CCI surgery (Figure 1f, g).

Furthermore, we detected the relationship of Syk-related proteins with FcγRI in DRG tissue after nerve injury. Coimmunoprecipitation (Co-IP) revealed the presence of Syk with FcγRI in the same complex of DRG tissue (Figure 1h). We found increased recruitment of Syk to FcγRI after nerve injury (Figure 1j). Further Co-IP showed that the proteins FcγRI, pSrc and Vav1 were related to the Syk protein and were more strongly bound to Syk proteins (Figure 1i, j). Immunofluorescence showed that Vav1, a downstream protein of the Syk signal, was also expressed in the DRG, colocalized with PGP9.5 (Figure 1k). Western blot results showed that CCI caused increased expression of phosphorylated Src (pSrc) in nerve-injured DRGs and that total Src did not significantly change in DRGs after CCI (Figure 1l, m). Similar to pSyk, pSrc was also mainly distributed in small neurons, and pSrc-immunopositive DRG neurons were more significantly increased in nerve-injured DRGs (Figure 1n).

### 3.3. Inhibition of neuronal FcγRI-Syk signaling mitigates chronic pain

CCI caused a substantial increase and activation of neuronal FcγRI/Syk signaling in the DRG. We continued to investigate the possible role in neuropathic pain. To determine the role of neuronal FcγRI in the development of nerve injury-induced neuropathic pain in the CCI model, we selectively deleted *Fcgr1* in DRG neurons by crossing rats carrying *loxP*-flanked *Fcgr1* with a primary sensory neuron– specific Cre line (*Pirt*^*Cre/+*^) to create *Fcgr1* conditional knockout (CKO) rats (Figure S1a-e). Compared with baseline, littermate control rats and CKO rats that underwent sham surgery did not show a decrease in the paw mechanical withdrawal threshold or thermal withdrawal latency (Figure S1f-g). However, the results of pain-related behaviors in CKO rats showed that both nerve injury-induced mechanical allodynia and thermal hyperalgesia were alleviated compared to littermate control rats (Figure 2a, b).

**Figure 2.**
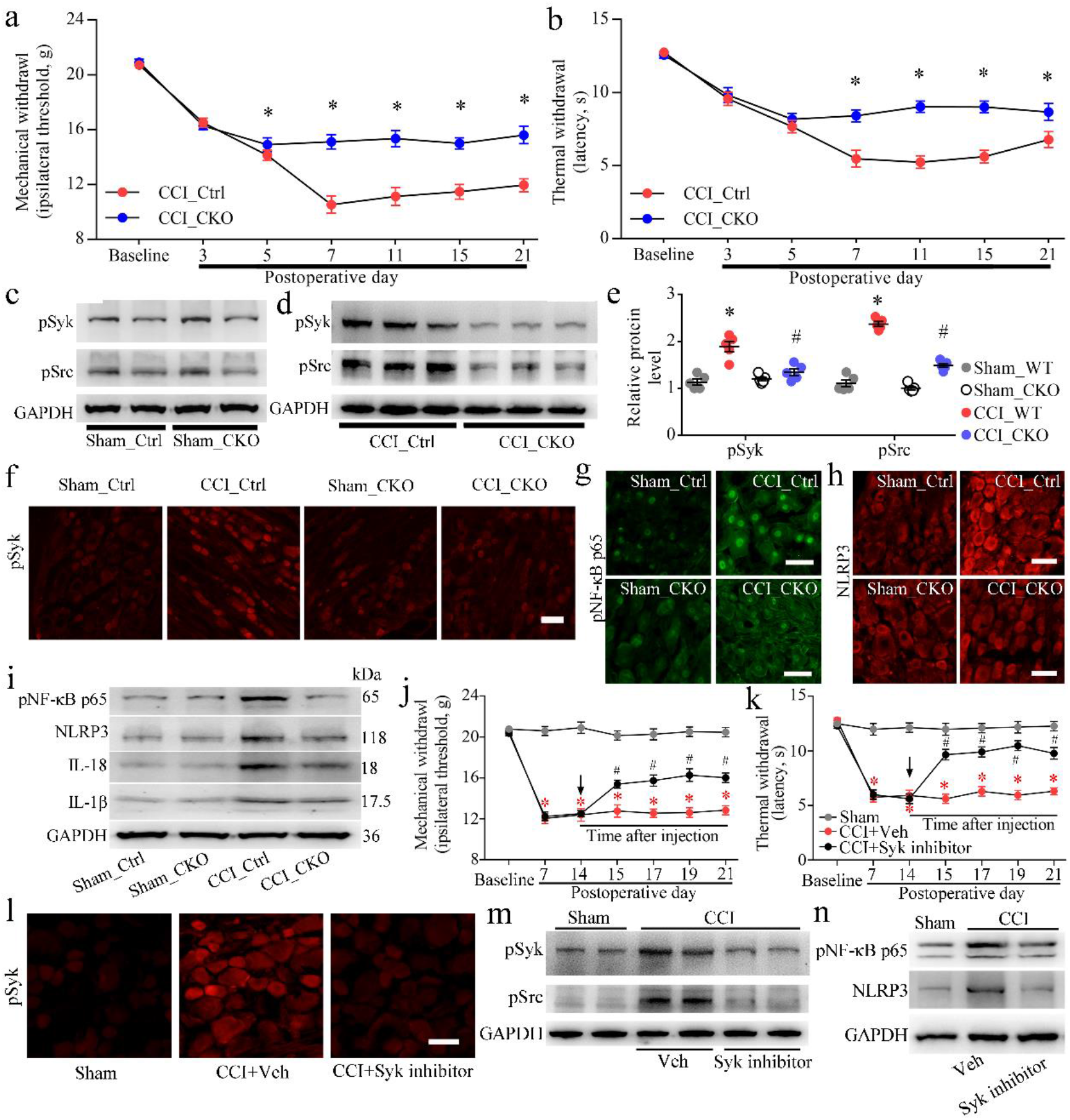
Neuronal FcγRI/Syk-related signaling activation mediated nerve injury-induced neuropathic pain. **(a-b)** Nerve injury-induced mechanical allodynia (c) and thermal hyperalgesia (d) were attenuated in *Fcgr1* CKO rats (n = 9/group). Two-way ANOVA, \**p* < 0.05, *Fcgr1* CKO vs. Ctrl. **(c-d)** Western blot showing the expression levels of pSyk and pSrc in Ctrl and *Fcgr1* CKO rat DRGs after sham (c) and CCI (d) injury. **(e)** Data summary of the expression levels of pSyk and pSrc in Ctrl and *Fcgr1* CKO rat DRGs after sham and CCI injury. **(f)** Immunofluorescence staining showing the pSyk cellular distribution in sham and CCI DRGs from Ctrl and CKO rats. **(g-h)** Immunofluorescence staining showing pNF-κB p65 (g) transport into the nucleus and NLRP3 (h) expression in sham and CCI DRGs from Ctrl rats and CKO rats 21 d after surgery. Scale bars: 50 μm in f, g and h. n = 4 rats/group. **(l)** Western blot showing the expression levels of pNF-κB p65, NLRP3, IL-18 and IL-1β in Ctrl and *Fcgr1* CKO rat DRGs after sham and CCI injury. **(j-k)** Nerve injury-induced mechanical allodynia (j) and thermal hyperalgesia (k) were attenuated in rats by a Syk inhibitor (n = 9/group). The first administration is indicated by an arrow on day 14 after CCI surgery. Rats were orally administered Syk inhibitor or vehicle b.i.d. from 14 d to 21 d after CCI surgery. Two-way ANOVA, \**p* < 0.05, CCI+Veh vs sham or CCI+Syk inhibitor. #*p* < 0.05, CCI+Syk inhibitor vs CCI+Veh or Sham. **(l)** Immunofluorescence staining showing the inhibitory effects of the Syk inhibitor on pSyk cellular distribution in DRGs 21 d after sham and CCI surgery. **(m)** Western blot showing the activation levels of pSyk and pSrc in DRGs from sham, CCI+Veh and CCI+Syk inhibitor rats 21 d after the operation. **(n)** Western blot showing the inhibitory effects of the Syk inhibitor on nerve injury-induced activation of pNF-κB p65 and NLRP3 in DRGs.

To directly determine whether Syk signaling pathways are involved in neuronal FcγRI-dependent neuropathic pain, we measured the effects of neuronal *Fcgr1* conditional knockout of DRGs on the phosphorylation of Syk and Src in CCI rats. Western blot results showed that the increased pSyk and pSrc levels in nerve-injured DRGs caused by CCI operation were significantly attenuated in the CKO rats after CCI (Figure 2c-e). Immunostaining showed that nerve injury-induced pSyk-immunopositive DRG neurons were significantly reduced in *Fcgr1* CKO rats (Figure 2f and Figure S1h). Further research found that phosphorylated NF-κB p65 (pNF-κB p65) was predominantly coexpressed with pSyk in DRG neurons after CCI (Figure S1i). In addition, immunostaining and Western blot results revealed that DRG neuron *Fcgr1* conditional deletion significantly reduced the nuclear transport and phosphorylation of NF-κB p65 and the expression of NLRP3 in DRG neurons (Figure 2g-i and Figure S1h, j and k). Furthermore, the nerve injury-induced upregulation of IL-1β and IL-18 was significantly decreased in the DRGs of CKO rats (Figure 1Sl, m).

In the CCI model of *Fcgr1* CKO rats, the activation of neuronal Syk signaling pathways was alleviated in the DRG. To further investigate the potential function of Syk signaling in nerve injury-induced neuropathic pain, we delivered a Syk inhibitor (PRT062607 (P505-15) HCl) or vehicle (normal saline) to rats on POD14 (orally twice a day). We found that Syk inhibitor administration daily for 7 consecutive days (from POD14 to POD21) produced long-lasting alleviation of nerve injury-induced mechanical allodynia and thermal hyperalgesia in rats (Figure 2j, k). To determine whether Syk/Src mediated neuronal FcγRI signaling, we detected the nerve injury-induced activation of Syk/Src signaling in the DRGs of CCI rats after Syk inhibitor administration. The immunostaining results revealed that nerve injury-induced phosphorylation of Syk (pSyk) in neurons were also alleviated in the DRGs after Syk inhibitor administration (Figure 2l). Meanwhile, Western blot results showed that nerve injury-induced pSyk, pSrc, pNF-κB p65 and NLRP3 expression in the DRG was significantly reduced after Syk inhibitor administration (Figure 2m, n).

### 3.4. Neuronal CRP induced pain and neuroinflammation via FcγRI

To investigate the cause of neuronal FcγRI/Syk signal activation after CCI, further experiments were performed to identify the neuronal FcγRI ligand. Previous studies found that C-reactive protein (CRP, also called PTX1) can bind and activate FcγRI ^23, 24^. Our western blot results revealed that nerve injury induced long-lasting increased expression of CRP (endogenous ligand of FcγRI) in the DRG (Figure 3a, b). To determine the cellular distribution of CRP, we then performed double immunolabelling using PGP9.5 with CRP. The results showed that CRP was more colocalized with PGP9.5 in DRGs after nerve injury (Figure 3c). To assess the behavioral effects of CRP, the intracutaneous injection of recombinant CRP was performed in the hindpaws of naïve rats. Intradermal injection of CRP induced dose-dependent mechanical and thermal hyperalgesia in naïve rats, whereas injection of vehicle (normal saline) did not evoke significant hyperalgesia compared with the baseline (Figure 3d, e). To investigate whether DRG neuronal FcγRI mediates CRP-induced pain, recombinant CRP protein was injected into the hindpaws of littermate control and CKO rats. The results revealed that CRP-induced mechanical and thermal hyperalgesia was significantly alleviated in CKO rats (Figure 3f, g). Previous research revealed that IgG-IC intracutaneous injection (1-10 μg/ml, 20 μl) can induce a significant decrease in the mechanical withdrawal threshold or thermal withdrawal latency ^14^. Further research found that recombinant CRP significantly enhanced IgG-IC intracutaneous injection-induced pain in rats (Figure 3h, i).

**Figure 3.**
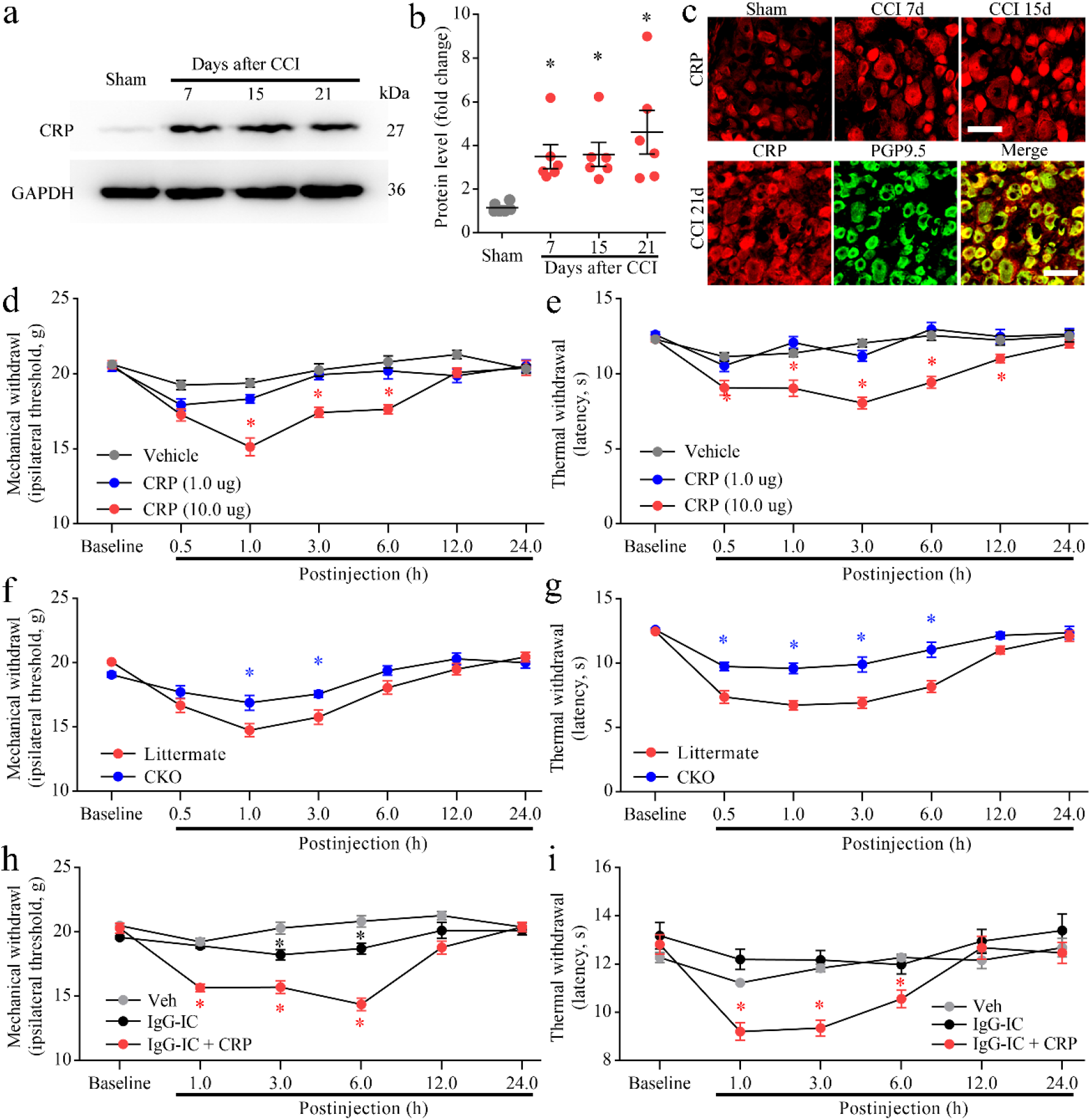
CCI increased CRP expression in the DRGs and CRP-induced pain in rats. **(a-b)** Western blot analysis (a) and quantification of CRP protein levels (b) in DRGs from sham, CCI 7 d, 15 d and 21 d rats after surgery. One-way ANOVA, n = 6 rats/group, \**p* < 0.05 versus the sham group. **(c)** Immunostaining of CRP in the DRGs of sham and CCI 7 d, 15 d and 21 d rats. Scale bars: 50 µm. n = 4 rats/group. **(d-e)** CRP induced mechanical (d) and thermal (e) hyperalgesia in naïve rats. Changes in the paw withdrawal mechanical threshold (d) and the paw withdrawal thermal latency (e) in the ipsilateral hindpaw after intradermal injection. Two-way ANOVA, \**p* < 0.05, vs Veh (PBS), n = 10 rats/group. **(f-g)** CRP-induced mechanical (f) and thermal (g) hyperalgesia was alleviated in CKO rats. Changes in the paw withdrawal mechanical threshold (f) and the paw withdrawal thermal latency (g) in the ipsilateral hindpaw after intradermal injection. Two-way ANOVA, \**p* < 0.05, vs littermate, n = 7 rats/group. **(h-i)** CRP sensitized IgG-IC-induced mechanical (h) and thermal (i) hyperalgesia in naïve rats. Changes in the paw withdrawal mechanical threshold (h) and the paw withdrawal thermal latency (i) in the ipsilateral hindpaw after intraplantar injection. Two-way ANOVA, \**p* < 0.05, vs Veh (PBS), n = 9 rats/group.

To further assess the pain-related function of CRP in DRG neurons, we overexpressed CRP in DRG neurons of rats by microinjection of the AAV2/9-Syn-CRP-ZsGreen vector into L4-L5 DRGs to encode CRP driven by the Syn promoter and establish a CRP overexpression model in vivo. Two weeks after injection of AAV2/9, CRP overexpression produced long-lasting mechanical and thermal hyperalgesia on the ipsilateral hind paw in littermate rats and *Fcgr1* CKO rats compared with naïve rats or AAV2/9-Syn-null-ZsGreen (Ad-Ctrl) injection rats (Figure 4a, b). Western blot results revealed that DRG injection of AAV2/9-CRP resulted in overexpression of CRP protein in the DRGs of littermate or CKO rats (Figure 4c, d). We then examined the effect of AAV2/9-Syn-CRP-ZsGreen (Ad-CRP-OV) on Syk signal activation. Compared with the Ad-Ctrl injection rats and naïve rats, the injection of Ad-CRP-OV induced activation of Syk (pSyk and pNF-κB p65) in the DRG of littermate rats. Further analysis showed that Ad-CRP-OV-induced increases in pSyk and pNF-κB p65 levels were significantly decreased in the Ad-CRP-OV injection group of CKO rats (Figure 4c, d).

**Figure 4.**
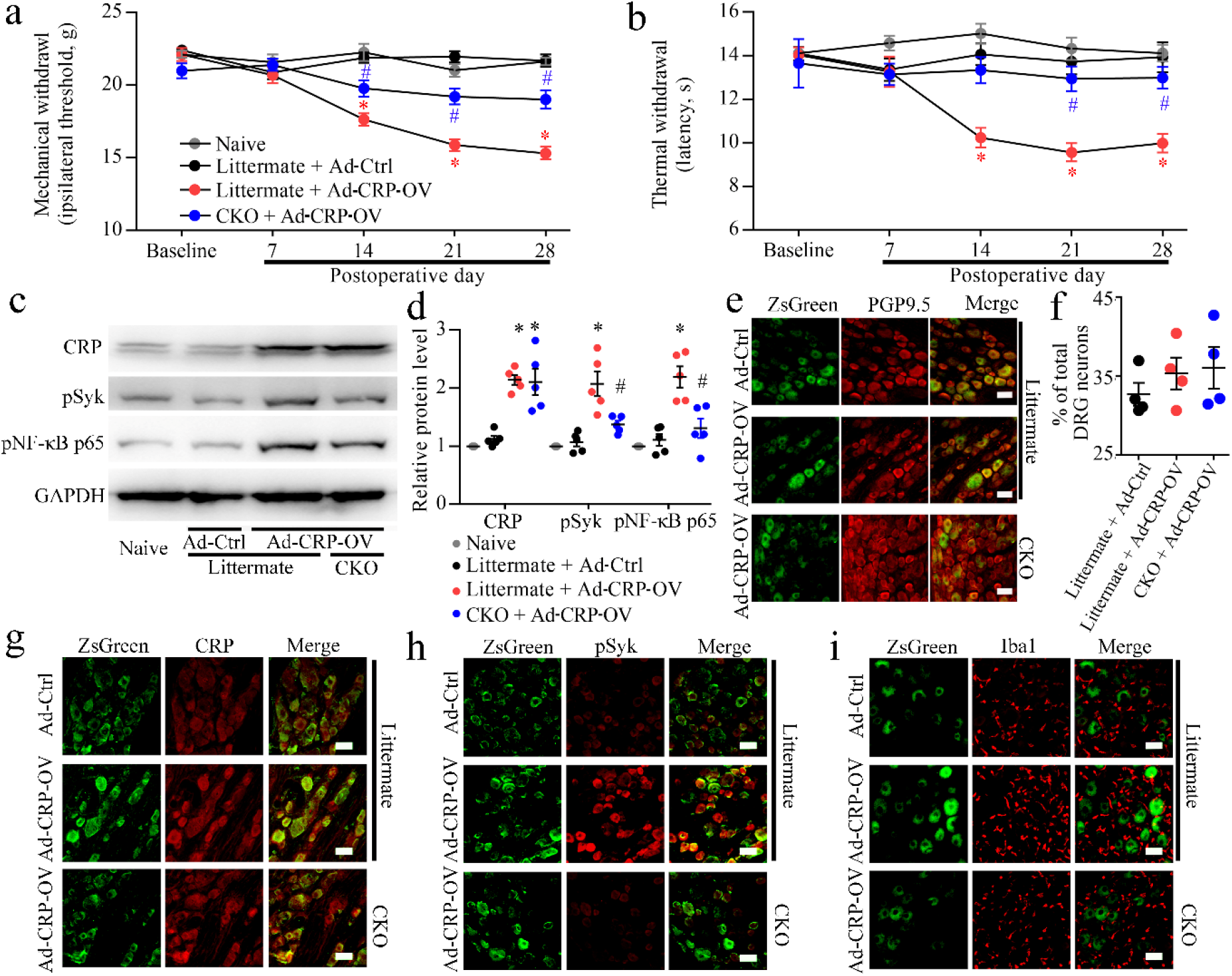
CRP overexpression induced pain and increased neuronal FcγRI-related neuroinflammation in DRGs. **(a-b)** CRP overexpression in the DRG induced FcγRI-mediated mechanical (a) and thermal hyperalgesia (b) in rats. Two-way ANOVA, n = 9 rats/group, \**p* < 0.05, Littermate + Ad-CRP-OV vs Naïve, Littermate + Ad-Ctrl or CKO + Ad-CRP-OV. #*p* < 0.05, CKO + Ad-CRP-OV vs littermate + Ad-CRP-OV. **(c)** Western blot showing the expression levels of CRP overexpression and activation of Syk/NF-κB p65 signals in DRGs of littermate and CKO rats after DRG injection of AAV2/9-hSyn-CRP. **(d)** Data summary of the expression levels of CRP, pSyk and pNF-κB p65 in the DRGs of littermate and CKO rats after DRG injection of AAV2/9-hSyn-CRP. One-way ANOVA, n = 5 rats/group, \**p* < 0.05 versus the naïve rats, #*p* < 0.05 versus the naïve group, littermate + Ad-Ctrl group, littermate + Ad-CRP-OV group. **(e)** Coexpression of the ZsGreen tag with PGP9.5 in neurons after DRG injection of AAV2/9-hSyn-CRP. **(f)** The positive rate of ZsGreen tag-positive neurons in the littermate + Ad-Ctrl group, littermate + Ad-CRP-OV group, and CKO + Ad-CRP-OV group. n = 4 rats/group. **(g)** Coexpression of the ZsGreen tag with CRP after DRG injection of AAV2/9-hSyn-CRP. **(h)** Coexpression of the ZsGreen tag with pSyk in neurons after DRG injection of AAV2/9-hSyn-CRP. **(i)** Coexpression of the ZsGreen tag with Iba1 after DRG injection of AAV2/9-hSyn-CRP. Scale bars: 50 µm in e, g, h, and i. Ad-CRP-OV = AAV2/9-hSyn-CRP-ZsGreen and Ad-Ctrl = AAV2/9-hSyn-null-ZsGreen in a–i.

Meanwhile, double immunolabelling showed that L_4_-L_5_ DRGs exhibited significant ZsGreen (green) labelling of neurons, resulting in an L_4_-L_5_ DRG neuron (PGP9.5-positive cells, red) labelling efficiency of 32.70% ± 1.45% in the Ad-Ctrl injection group of littermate rats, 35.33% ± 2.02% in the Ad-CRP-OV injection group of littermate rats, and 36.04% ± 2.64% in the Ad-CRP-OV injection group of CKO rats (Figure 4e, f). Immunolabelling images showed that ZsGreen labelling of DRG neurons also expressed CRP after Ad-CRP-OV injection (Figure 4g). Compared to Ad-Ctrl injection, pSyk-positive neurons and Iba1-positive cell infiltration in the DRG were increased after injection of Ad-CRP-OV in littermate rats and significantly reduced in the DRG of CKO rats (Figure 4h, i). These findings indicate that blocking neuronal FcγRI activation in DRGs can suppress CRP-dependent chronic pain in a rat CRP overexpression model.

### 3.5 DRG injection of native IgG suppresses persistent pain and neuroinflammation after CCI

In CRP-overexpressing rats, which displayed enhanced hyperalgesia by CRP, we next performed DRG application of native IgG to investigate the effect of analgesia in the CCI model of rats. The application of native IgG gradually increased the nociceptive (mechanical and thermal) threshold evoked by the CCI operation (Figure 5a, b). To evaluate the effects of native IgG on gene expression in nerve-injured DRGs, RNA sequencing was used to analyse the messenger RNA (mRNA) profiles in DRGs obtained from rats with sham operation, nerve injury (vehicle application) or nerve injury (native IgG application). Compared with the sham group, the nerve injury group treated with vehicle had 473 differentially expressed genes (DEGs) in the injured DRGs (Figure 5c). A total of 361 DEGs, 105 upregulated and 256 downregulated, were identified in the nerve-injured DRGs with native IgG application compared with the nerve-injured DRG group treated with vehicle (Figure 5d). A heatmap of the mRNA expression levels in the DRGs of the three groups (sham operation, CCI injury with vehicle and CCI injury with native IgG) is shown in Figure 5e. We comparatively analysed the overlapping mRNA of DEGs in nerve injury with vehicle group vs. sham group and nerve injury with native IgG group vs. nerve injury with vehicle group DRGs. A Venn diagram was presented to depict the overlaps between the two sets of DEGs (Figure 5f). We further identified multiple signaling pathways relevant to both inflammation and immune regulation, such as the FcγR signaling pathway, cytokine–cytokine receptor interaction, chemokine signaling pathway, and Toll-like receptor signaling pathway, by KEGG analysis (Figure 5g).

**Figure 5.**
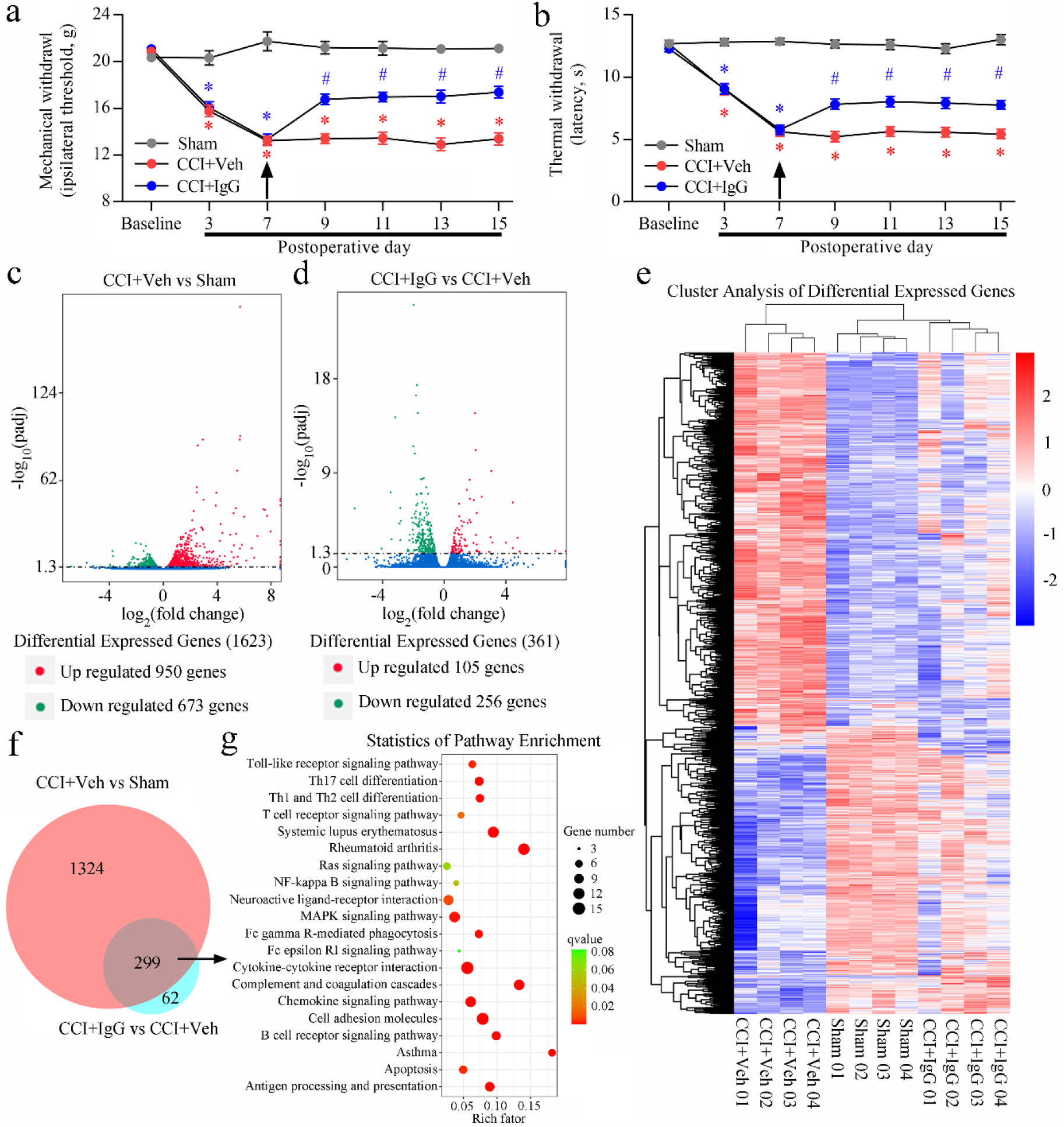
Inhibitory effects of native IgG on nerve injury-induced pain hypersensitivity after DRG injection. **(a-b)** Nerve injury-induced mechanical allodynia (a) and thermal hyperalgesia (b) were attenuated in rats by native IgG. Two-way ANOVA, n = 9/group, *p < 0.05, CCI vs sham or CCI + IgG. #p < 0.05, CCI + IgG vs CCI. **(c)** Volcano plot showing the overall distribution of upregulated and downregulated mRNAs between the sham group and CCI+Veh group rats (n = 4 per group). **(d)** Volcano plot showing the overall distribution of upregulated and downregulated mRNAs between the CCI+Veh group and CCI+IgG group (n = 4 per group). **(e)** Heatmap of mRNAs showing hierarchical clustering of differentially expressed mRNAs in the 3 groups. In the clustering analysis, upregulated genes (red) and downregulated genes (blue) are shown. **(f)** Venn diagram indicating the numbers of overlapping and distinct mRNAs between the CCI+Veh vs sham group and CCI+IgG vs CCI+Veh group. **(g)** KEGG pathway scatterplot of genes associated with overlapping mRNAs from f. Rich factor, qvalue, and Gene number of enriched in this pathway were used to measure the enrichment degree of genes in the KEGG analysis. Rich factor refers to the number of differentially expressed genes enriched in a pathway as a ratio of the number of annotated genes. The rich factor value represents the enrichment degree. The qvalue represents a p value corrected by multiple hypothesis tests.

mRNA sequencing revealed that the FPKM values of *Fcgr1* and related proinflammatory cytokine genes in DRGs were increased after nerve injury and significantly decreased after native IgG microinjection (Figure 6a). Here, we further explored whether native IgG attenuated neuropathic pain by suppressing neuronal FcγRI/Syk signal activation. We also examined the changes in NF-κB p65-related proinflammatory signaling. Western blotting found that DRG microinjection of native IgG significantly reduced the expression level of FcγRI and phosphorylation level of Syk (pSyk), which was induced by CCI (Figure 6b, c).

**Figure 6.**
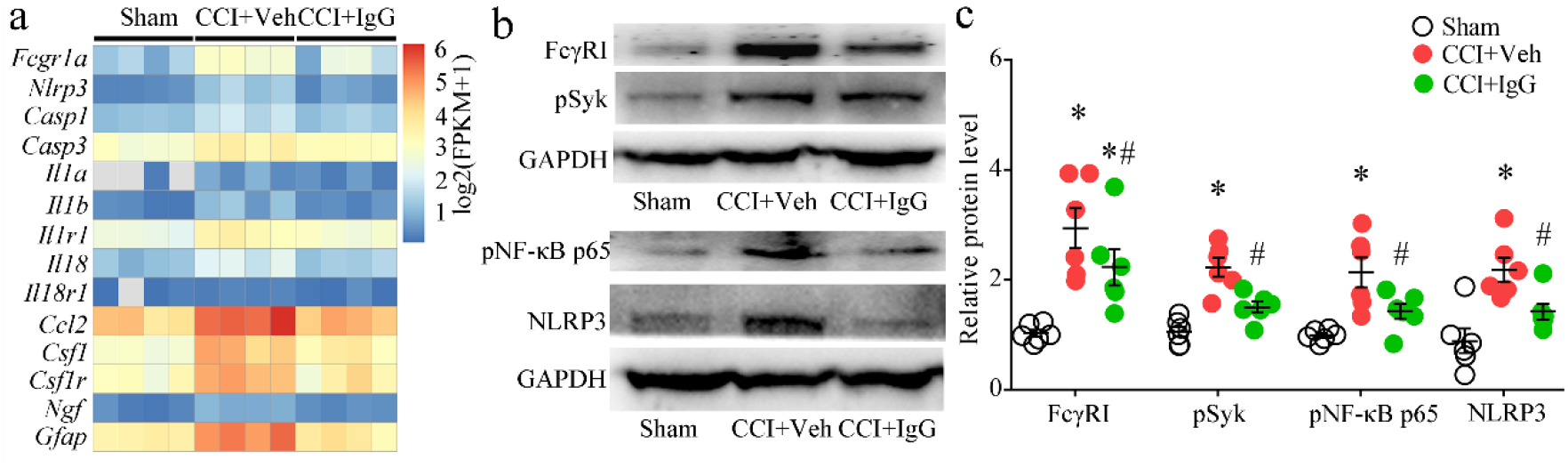
Native IgG suppresses nerve injury-induced FcγRI/pSyk/NLRP3 signaling activation after DRG injection. **(a)** The heatmap shows the differential expression of *Fcgr1* and proinflammatory cytokine genes in the DRGs of sham, CCI+Veh and CCI+IgG rats. **(b)** Western blot analysis of the effects of native IgG on the neuronal FcγRI/pSyk/pNF-κB p65/NLRP3 signaling pathway in DRGs after CCI injury. **(c)** Data summary of the protein expression levels of FcγRI, pSyk, pNF-κB p65 and NLRP3 in the inhibitory effects of native IgG on nerve injury-induced rat DRGs after operation. n = 6 rats/group. \**p* < 0.05 versus the sham group, #*p* < 0.05 versus the CCI+Veh group, by one-way ANOVA.

## Discussion

Our study reveals a critical role of neuronal CRP/FcγRI/Syk signaling in the neuropathic pain of nerve injury. Nerve injury-induced neuronal FcγRI-related Syk signaling activation may contribute to neuropathic pain after nerve injury by regulating the proinflammatory cytokines IL-1β and IL-18 in neurons in the DRG. Nerve injury also causes rapid and long-lasting high expression of CRP in DRG neurons. CRP may be involved in the production and persistence of chronic pain in the CCI model of rats by activating the neuronal FcγRI-related Syk signaling pathways of DRGs. DRG conditional knockout or DRG native IgG blockade of neuronal FcγRI alleviates nerve injury-induced neuropathic pain. Meanwhile, inhibition of neuronal FcγRI suppresses nerve injury-induced activation of the Syk/NF-κB p65 proinflammatory signaling pathway and induction of neuronal IL-1β and IL-18 in the DRG, as well as activation of astrocytes and microglial cells in the spinal dorsal horn. The results of our present study further confirm that FcγRI activation of sensory neurons may contribute to neuropathic pain. These findings may support a new understanding of the mechanism and therapy of neuropathic pain after nerve injury.

FcγRs belong to the immunoglobulin superfamily and are classified as activating or inhibitory. In humans, FcγRs include hFcγRI, FcγRIIA, hFcγRIIB, hFcγRIIC, hFcγRIIIA, and hFcγRIIIB. Mice express four FcγRs (I-IV), and rats express three FcγRs (I-III) ^25-28^. Activating FcγRs contain immunoreceptor tyrosine activating motifs (ITAMs), and inhibitory FcγRs contain immunoreceptor tyrosine inhibitory motifs (ITIMs) ^27, 28^. FcγRI is known to be expressed most prominently on immune cells, including macrophages, dendritic cells, natural killer cells, neutrophils, eosinophils, and mast cells ^26-28^. Beyond immune cells, recent studies have shown that neurons, astrocytes, and microglial cells of the central nervous system (CNS) and peripheral nervous system (PNS) also express FcγRs, including FcγRI ^8, 9, 29-32^. In addition, sensory neurons of the DRG express FcγRI, as reported by several studies ^8, 9, 11, 12^. Previous studies found that FcγRI in DRGs also contributes to joint pain in rodent models of RA. In autoimmune diseases such as RA, FcγRI, which is expressed on nociceptors of the DRG, is activated by autoantigen-antibody-IC ^19^. Autoantibodies have also been found to be a potential mechanism driving pain in complex regional pain syndrome (CRPS). Autoantibodies against voltage-gated potassium channels (VGKCs) can be detected in patient blood samples of CRPS. Many studies have found that autoimmunity plays a role in the initiation and maintenance of neuropathic pain ^5, 7^. Depletion of B cells may also be a disease-modifying treatment for pain in a rodent model of CRPS ^33^. We showed that intraplantar injection of IgG-IC rapidly induced mechanical and thermal hyperalgesia in WT rats.

FcγRI is the only high-affinity activating FcγR and binds to the Fc portion of IgG. Multimeric IgG-IC crosslinks FcγRI to enable receptor clustering, aggregation, and activation, leading to ITAM domain phosphorylation. The recruitment and activation of Src and Syk kinases are crucial steps in the activation of FcγRI ^34-36^. Our previous study demonstrated that IgG-IC activates neuronal FcγRI in DRGs by coupling with the TRPC3 cation channel through the Syk signaling pathway ^22^. In a rat model of RA, the phosphorylation levels of Src and Syk were increased in DRG neurons ^10^. In this study, several key molecules (Syk, Src and NF-κB p65) in the FcγRI signaling pathway were found to be significantly regulated in DRG neurons of rats after nerve injury. In addition to the interaction of FcγRI with IgG in passive immunization, FcγRI is also activated by CRP, which is an innate immunity-related protein ^37-39^. The innate immune response is the first line of defense against sterile tissue damage and infection ^40, 41^.

CRP is a member of the pentraxin family, a ligand for Fc receptors on phagocytes and a major acute phase protein in tissue damage and inflammation ^23, 42, 43^. CRP is unanimously regarded as an inflammatory biomarker associated with depression, schizophrenia, posttraumatic stress disorder and autism ^44-48^. Previous studies have reported that CRP has a direct proinflammatory effect on endothelial cells and enhances IgG-mediated phagocyte responses in immune thrombocytopenia ^49, 50^. CRP can induce the expression of pro-IL-1β and NLRP3 and activate the NLRP3 inflammasome in endothelial cells via the FcγR/NF-kB pathway ^51^. In the present experiments, our results showed that neuronal CRP in the DRG was increased after nerve injury, and intraplantar injection of CRP produced mechanical and thermal hyperalgesia and aggravated IC-induced pain in rats. Previous studies on the chronic pain of arthritis have reported that conditional deletion of *Fcgr1* in sensory neurons significantly reduces IgG-IC-induced pain in arthritis models in mice and rats ^10, 19^. We observed that deletion of *Fcgr1* in sensory neurons of the DRG significantly alleviated CRP-induced pain in *Fcgr1* CKO rats. CRP overexpression in DRG-induced pain was also reduced in *Fcgr1* CKO rats.

Intravenous immunoglobulin (IVIG) is widely used in the immunotherapy of autoimmune and inflammatory diseases such as RA, Kawasaki disease, systemic sclerosis, Guillain–Barre syndrome, and chronic inflammatory demyelinating polyneuropathy ^52^. IVIG has been shown to be effective in treating inflammation-related pain symptoms of rheumatic diseases. Previous research has revealed that IVIG may be beneficial for managing neuropathic pain ^53^. Some research has reported that IVIG and plasma exchange therapy could effectively improve pain symptoms in some CRPS patients ^54-56^. We examined the therapeutic effect of IgG injection in the local DRG and found persistent pain relief in CCI rats. Native IgG Fc alone has anti-inflammatory or immunomodulatory activity ^24, 57-59^. A high level of monomeric IgG can bind FcγRI and block or reduce the IC-mediated activation of FcγRI ^36, 57, 58^. Based on nociceptive neurons of the DRG expressing FcγRI, we found that IgG injection of local DRG reduced the local neuroinflammation of DRG by modulating neuronal FcγRI/Syk signaling.

## Conclusions

The present study supports a novel view of how nerve injury-induced CRP mediates the nociceptive behavior of neuropathic pain. Local formation of native IgG has the potential to serve important roles in the therapy of chronic pain. Taken together, these results suggest that neuronal FcγRI-related signaling plays an important role in neuropathic pain. These findings may provide novel insights into the interactions between nerve damage and peripheral neuroimmunity in pathologic conditions.

## Authors’ contributions

F.L. and C.M. designed the experiments and wrote the manuscript. F.L., S.S. and L.Z. performed the experiments and analysed the results. Y.F., H.C., J.S. and S.J. acquired and analysed the data.

## Data Sharing statement

All data will be made available by the corresponding author upon reasonable request.

## Study approval

All animal studies were approved by the Institutional Animal Care and Use Committee of the Institute of Basic Medical Sciences, Chinese Academy of Medical Sciences, Peking Union Medical College (Beijing, China).

## Acknowledgements

We thank Dr. Wenyin Qiu, Dr. Xiaojin Qian, Dr. Yongmei Chen, Dr. Tao Wang and Ms. Bo Yuan (Department of Anatomy, Histology and Embryology, Institute of Basic Medical Sciences, Chinese Academy of Medical Sciences) for technical assistance in immunohistochemistry.

## Supplementary Methods

### Rat Genotyping

The *Fcgr1* CKO (*Fcgr1*^*loxP+/+*^;*Pirt*^*CRE*+^) rat genotype was identified by PCR. Genomic DNA was prepared from 5 mm clipped tail samples from rats and extracted in DNA lysis buffer containing proteinase K using the Tail Genomic DNA Kit (Cat: CW2049S, CoWin Biosciences). The DNA solution of the rat genotype was identified by PCR with 2×Taq MasterMix (Cat: CW0682S, CoWin Biosciences). DNA was predenaturated at 94°C for 2 mins, followed by PCR with 30 cycles of 94°C for 30 s, 50°C for 30 s, and 72°C for 30 s. The primers for *Fcgr1loxP* were as follows: forward, 5′-CTGTAATTCTGCTACTGTTATGAATCGTAATG-3′ and reverse, 5′-CCTGAACACCAAAATCTGCCTG-3′. The primers for *Pirt-Cre* were as follows: forward, 5′-TACTGACGGTGGGAGAATG-3′ and reverse, 5′-CTGTTTCACTATCCAGGTTACG-3′.

### Fluorescent in situ hybridization

To examine the expression of *Fcgr1* mRNA in DRG neurons of rats, fluorescent in situ hybridization (FISH) was used with locked nucleic acid probes specific for *Fcgr1*. Rats were sacrificed under anesthesia. The harvested DRGs were fixed with 4% paraformaldehyde. Protease K (20 μg/ml) was added to digest the frozen DRG sections at 37°C for 5 minutes. After washing with pure water, the sections were washed 3 times for 5 min each with PBS. After preincubation in hybridization solution at 37°C for 1 h, the sections were incubated overnight in hybridization solution with 8 ng/μl probes at 37°C. DAPI dye was added to the sections and incubated in the dark for 8 min. After rinsing, anti-fluorescence quenching sealant was added to the sections, which were then sealed. The sequence of *Fcgr1* probes: 5’Cy3-TTGCCCACCAACTGGAACCCAAAGTA-3’, the sequence of *Fcgr1* exon 3 mRNA: 5’FITC-CACCACUGUCCUUGAAACUGGCCUUGAGGAUGC-3’.

## Supplementary Data

**Figure S1.**
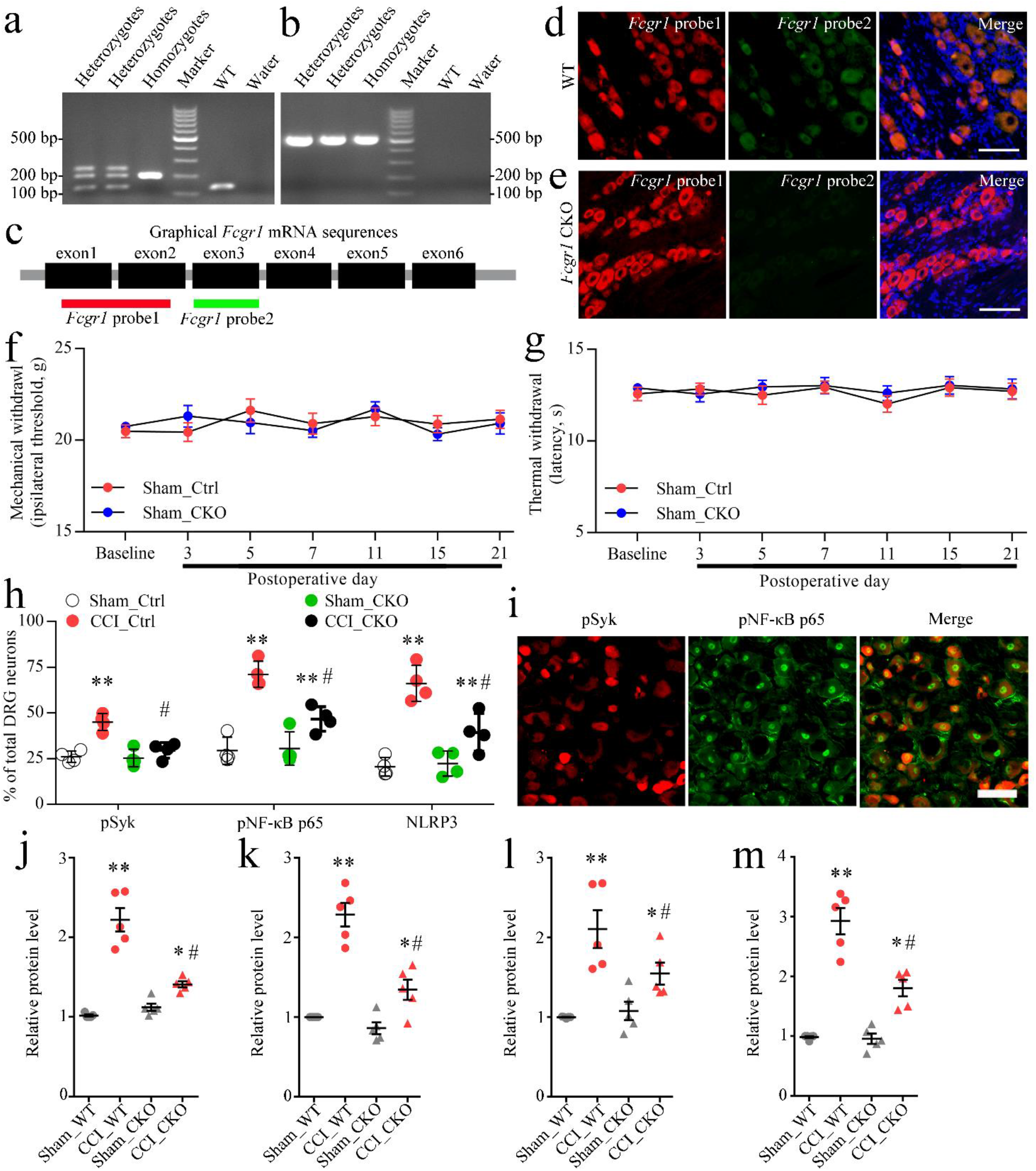
**(a)** PCR analysis of the genotypes *loxP-flanked Fcgr1* in the CKO rats and WT rats. **(b)** RT–PCR analysis of *Pirt-Cre* expression in CKO rats and WT rats. **(c)** Diagram showing *Fcgr1* mRNA targets identified with fluorescence in situ hybridization (FISH) probe 1 (red) and FISH probe 2 (green). **(d-e)** FISH analysis using the *Fcgr1* probe, revealing *Fcgr1* mRNA exon 3 expression (green) on DRGs from WT rats (d) and CKO rats (e). Homozygotes: *Fcgr1*^*loxP+/+*^; *Pirt*^*CRE*+^ (*Fcgr1* CKO) rats, Heterozygotes: *Fcgr1*^*loxP+/-*^; *Pirt*^*CRE*+^ rats, WT: wild-type rats. **(f-g)** Comparison of the von Frey (a) and heat (b) withdrawal thresholds between *Fcgr1* CKO and littermate control (Ctrl) rats after sham operation, n = 9/group. **(h)** The positive rate of pSyk, pNF-κB p65, and NLRP3 in sham and CCI DRGs from littermate control (Ctrl) and CKO rats 21 d after surgery. n = 4 rats/group. \*\**p* < 0.01 versus the Sham_Ctrl group and Sham_CKO group, #*p* < 0.05 versus the CCI_Ctrl group, by one-way ANOVA. **(i)** Immunofluorescence staining showing cellular colocalization of pSyk (red) and pNF-κB p65 (green) in the DRGs of rats 21 d after CCI surgery. **(j-m)** Data summary of the protein expression levels of pNF-κB p65 (j), NLRP3 (k), IL-18 (l) and IL-1β (m) in sham and CCI DRGs from Ctrl and CKO rats 21 d after surgery. n = 5 rats/group. \**p* < 0.05 versus the Sham_Ctrl group, #*p* < 0.05 versus the CCI_Ctrl group, by one-way ANOVA.

